# An Efficient Strategy to Estimate Thermodynamics and Kinetics of G Protein-Coupled Receptor Activation Using Metadynamics and Maximum Caliber

**DOI:** 10.1101/367888

**Authors:** Derya Meral, Davide Provasi, Marta Filizola

## Abstract

Computational strategies aimed at unveiling the thermodynamic and kinetic properties of G Protein-Coupled Receptor (GPCR) activation require extensive molecular dynamics simulations of the receptor embedded in an explicit lipid-water environment. A possible method for efficiently sampling the conformational space of such a complex system is metadynamics (MetaD) with path collective variables (CV). Here, we applied well-tempered MetaD with path CVs to one of the few GPCRs for which both inactive and fully active experimental structures are available, the μ-opioid receptor (MOR), and assessed the ability of this enhanced sampling method to estimate thermodynamic properties of receptor activation in line with those obtained by more computationally expensive adaptive sampling protocols. While n-body information theory (nBIT) analysis of these simulations confirmed that MetaD can efficiently characterize ligand-induced allosteric communication across the receptor, standard MetaD cannot be used directly to derive kinetic rates because transitions are accelerated by a bias potential. Applying the principle of Maximum Caliber (MaxCal) to the free-energy landscape of morphine-bound MOR reconstructed from MetaD, we obtained Markov State Models (MSMs) that yield kinetic rates of MOR activation in agreement with those obtained by adaptive sampling. Taken together, these results suggest that the MetaD-MaxCal combination creates an efficient strategy for estimating thermodynamic and kinetic properties of GPCR activation at an affordable computational cost.

## I. INTRODUCTION

G protein-coupled receptors (GPCRs) are broadly expressed cell surface receptors whose functional role is to transmit signals from the exterior to the interior of the cell through recognition of different ligands, such as bioactive peptides, amines, nucleosides, and lipids.^1, 2^ It is therefore not surprising that about 30% of drugs available in the market today target GPCRs for the purpose of alleviating the effects of a wide range of diseases and conditions.^3^ Understanding the mechanistic, thermodynamic, and kinetic details of ligand-induced GPCR activation and consequent signaling through intracellular G proteins or β-arrestins is very important as it informs the rational design of improved therapeutics. However, this remains a fairly challenging undertaking both for experimental and computational approaches.

Molecular dynamics (MD) simulations are a particularly valuable tool to probe the level of atomistic detail that is necessary to identify testable hypotheses of molecular determinants that are responsible for GPCR allostery, energetics, and kinetics. However, simulating ligand-induced activation of GPCRs in a realistic 1 lipid-water environment is computationally expensive and one cannot rely on standard MD alone to obtain converged free-energy landscapes, as the largest conformational changes accompanying receptor activation occur at the timescale of hundreds of microseconds to milliseconds.^4^

Enhanced sampling algorithms can help speed up these timescales as we demonstrated a few years ago by applying well-tempered metadynamics^5^ (MetaD) with path collective variables (CVs)^6^ to study the ligand-induced modulation of the free-energy landscape of two prototypic GPCRs, specifically rhodopsin^7^ and β2-adrenergic^8^ receptors embedded in an explicit 1-palmitoyl-2-oleoyl-sn-glycero-3-phosphocholine (POPC)/10% cholesterol bilayer. More recently, using a high-throughput molecular dynamics (HTMD) adaptive sampling protocol^9, 10^ run on large distributed computational resources, we demonstrated^11^ that without introducing bias potentials a total simulation time of ∼240 µs was still necessary to achieve convergence of the free-energy landscape of a morphine-bound μ-opioid receptor (MOR) system embedded in an explicit POPC/10% cholesterol bilayer, and to build reliable Markov State Models (MSMs) of the system’s activation dynamics. This approach successfully revealed the presence of two distinct metastable regions of the conformational space in addition to metastable regions comprising crystal-like inactive and active conformations of MOR. Furthermore, it unveiled ligand-specific kinetic rates between these regions and, combined with information theory,^12^ it elucidated molecular details of the allosteric transmission of the signal across the receptor. However, the conformational sampling portion of this approach is still computationally very expensive and often requires exclusive computational resources.

Here, we propose a computational strategy that combines well-tempered MetaD using path CVs^6^ with Maximum Caliber (MaxCal) and n-body information theory^13^ (nBIT) to efficiently study ligand-induced activation of GPCRs at the atomistic scale. Using MetaD-derived free energies as an input for Maximum Caliber (MaxCal)^14-16^, we built MSMs and estimated kinetic rates of morphine-induced MOR activation. Moreover, the n-body information theory method nBIT was used to elucidate molecular details of MOR allosteric modulation leading to signal transmission. The efficiency and accuracy of the proposed approach was evaluated by comparing these results with those obtained with a more expensive HTMD adaptive sampling protocol run on distributed computational resources.^11^

## II. COMPUTATIONAL DETAILS

### A. Setup of ligand-free active and inactive MOR systems

Inactive and active three-dimensional models of the MOR were built based on corresponding available experimental structures at the start of this work (PDB 4DKL^17^ and 5C1M^18^, respectively). For the active system, the nanobody was removed, the N-terminal region was truncated at residue S64^1.40^, and the missing residues of helix 8 (H8) were modeled using as a template the atomic coordinates of H8 in the inactive crystal structure. The receptor termini were capped with an N-terminal acetyl and a C-terminal N-methyl amide using Maestro. An 80×80 Å^2^ POPC/10% cholesterol bilayer with TIP3P water was generated using the CHARMM-GUI webserver^19^ and equilibrated according to its standard protocol, comprising a 20 ns unrestrained MD simulation of the membrane. The *inflategro* script,^20^ edited to use a deflation ratio of 0.97 instead of the default 0.95, was used to embed the final MOR structures in the membrane mimetic environment. After system neutralization with NaCl at a concentration of ∼150 mM, the entire simulation system of approximately 54,000 atoms with dimensions of 75×75×100 Å^3^ was minimized and equilibrated for 50 ns using the CHARMM36 force field^21, 22^ for protein, lipids and ions. All simulations reported in this work were carried out using the GROMACS 5.1.4^23^ package.

### B. Setup of morphine-bound active MOR system

Force-field parameters for morphine were obtained from the CHARMM General Force Field (CGenFF) ParamChem webserver^24^ and subsequently optimized and verified following established protocols^24^. The initial binding pose of morphine was obtained by aligning the ligand’s alkaloid scaffold with the equivalent one in the morphinan agonist BU72 as seen in the 5C1M crystal structure.^25^ The system was subjected to minimization followed by a 1 ns run in the NVT ensemble with restraints on lipids, receptor, and ligands. Restraints were gradually relaxed over 20 ns and a final 40 ns equilibration run was carried out without restraints.

All production runs were carried out in the NPT ensemble at 300 K and 1 bar with periodic boundary conditions. The Parinello-Rahman^26^ and velocity rescaling^27^ algorithms were used for pressure and temperature coupling, respectively. A time step of 4.0 fs coupled with hydrogen mass repartitioning was used alongside the standard leapfrog algorithm,^28^ and the LINCS algorithm.^29^ Electrostatic interactions were handled via the Particle-Mesh Ewald method^30^ and a Verlet scheme^31^ with a cutoff of 1.2 nm, while the van der Waals modifier force switch^32, 33^ was set to 1.0 nm.

### C. Adiabatic biased MD simulations

To obtain an initial representation of the transition between the inactive and active crystal structures of MOR, we carried out adiabatic biased MD (ABMD) simulations^34^ on the ligand-free receptor using Plumed 2.1.^35^ Specifically, 10 simulations were carried out starting from the equilibrated active MOR crystal structure guided towards the receptor inactive conformation, and 10 simulations were run in the opposite direction. In ABMD simulations, the system is biased with an elastic potential towards a final value of a chosen CV. The bias, however, acts on the system only when the distance of the current value of the CV to its final value is larger than its previous minimum distance, allowing the system to evolve undisturbed otherwise. The difference between the contact map calculated during the simulation (with a stride of 10 simulation steps) and the contact map of the final structure (i.e., that of the inactive or active MOR structures, depending on the direction of the simulation), defined by a subset of contacts relevant to the activation process, was used as a CV. Specifically, a switching function in the form of

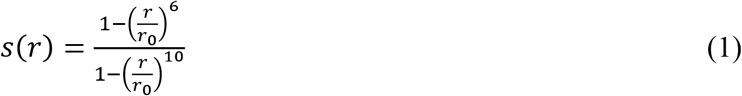

was used to describe contacts between polar atoms or the side chains of residues within the MOR transmembrane region. In this equation, *r* is the distance between two polar atoms or between the center of mass of apolar side chains, and *r*_0_ was set to 6.5 Å or 4.5 Å for side chain contacts or polar contacts, respectively. Contacts for which |*s*(*r*_act_) − *s*(*r*_inact_)| > 0.65, i.e. whose distance in the active and inactive structures are significantly different, were included in the contact map definition *R*_*ij*_ = *s*(*r*_*ij*_), and the matrix norm ‖*R*^Ref^ − *R*‖, where *R*^Ref^ is either the contact map for the active or for the inactive receptor structure, was used to drive the ABMD simulations. The simulations were performed in a stepwise fashion with the force constant switched from 0.1 to 15 over the span of 35 ns to ensure a smooth transition between receptor conformations.

### D. Path definition for MetaD simulations

In order to define a path based on information from contact maps, a pairwise distance matrix ‖*R_k_* − *R_k′_*‖ was built for the complete set of frames from the ABMD trajectories by running an in-house python script on contact maps calculated with the Plumed 2.1^35^ plugin. In order to select frames optimally describing the low free-energy de-/activation pathway, we performed simulated annealing over paths {j_m_} of constant length *N* = 10 starting from a random initial selection of 8 points between the two furthest points in the set, j_0_ and j_*N*_, respectively, and minimizing the function

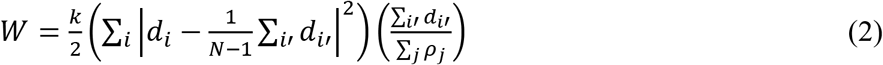

where *i, i*′ enumerate edges along the path, *d*_*i*_ = ‖*R(j_*i*_)* − *R*(*j*_*i*+1_)‖ is the length of the *i*-th edge, ρ_j_ represents the density of neighboring points around the beginning point *j*, and *k* is the effective temperature for the annealing procedure, which was slowly decreased over the span of the annealing iterations. Minimization of this function, which was performed using an in-house python script, allows us to (a) find the most densely populated regions along the activation path, (b) ensure that each edge is of similar length, and (c) verify that the overall length of the path is the shortest possible while fulfilling the two prior conditions. The value of *W* converged to yield paths with edge length variances below 0.05, ensuring that the chosen frames could be used to define smooth path collective variables that describe the position *S*(R) of the system along the pathway, and the distance *Z*(R) from this path. These were defined, respectively, as:

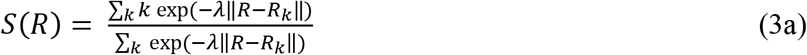

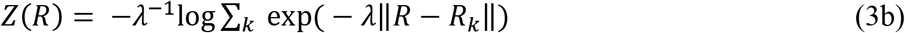

where R_k_ refers to the values of the contact maps for each point in the selected path, and λ, a parameter to aid the creation of a smooth description of the path, is set to ∼2.0.

### E. MetaD Simulations

We used well-tempered MetaD with the path collective variables *S* and *Z*, as implemented in Plumed 2.1^35^, to enhance the exploration of the CV space by adding a history-dependent bias potential at regular intervals to discourage the system from visiting previously explored regions of the conformational space. Specifically, the bias potential acting on a conformation with contact map R is

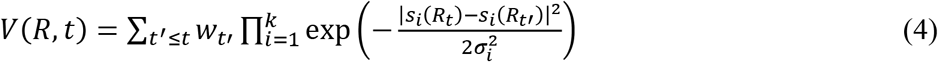

where *t*′ is a multiple of the deposition time τ, *s*_*i*_ are the CVs over which the bias is being deployed (*S* and *Z* of the path CVs in our case), and the σ_*i*_ are the standard deviations of the Gaussian bias. In well-tempered MetaD the height of the Gaussian bias, *w*_*t*′_, is adjusted according to

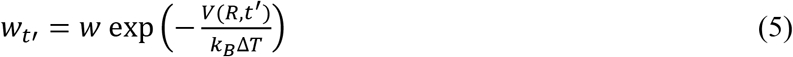

where Δ*T* is a parameter in the units of temperature, k_*B*_ is the Boltzmann constant, and *w* is the initial height of the Gaussian bias potential. This adjustment allows the total bias potential to smoothly converge in time so that the free energy of the system can be calculated as the limit:

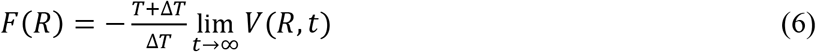

where *T* is the temperature of the simulation. The temperature ratio Δ*T*/(*T*+Δ*T*) in equation 6 is referred to as the “bias factor” and ensures that the relevant free-energy barriers can be overcome within the timescale of metadynamics simulations. Here, we run two sets of simulations of the morphine-bound MOR system, where the bias factor was set to 12, the deposition rate to τ = 5 ps, and either (σ_*S*_ = 0.3, σ_*Z*_ = 0.15) or (σ_*S*_ = 0.1, σ_*Z*_ = 0.05) were used. Both simulations were run for 1.5 μs. Convergence was checked by ensuring that the standard deviation of the free-energy differences converged to below 20 kJ/mol within the last 150 ns for both simulations. We present the combined free energies and the standard deviations of the free-energy differences in Supplementary Fig. 1(a) and 1(b), respectively, where the two simulations, totaling 3 µs, were combined using weights of 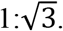

### F. Free-Energy Calculations

To derive distributions of order parameters other than the path collective variables, it is necessary to remove the effect of the bias on the trajectories of sampled conformations. This can be achieved using the reweighting method introduced by Tiwary et al.^36^ and implemented via in-house python scripts, which allows to reconstruct a time-independent free-energy landscape for any function of the coordinates of the system. Specifically, this method was used to recast the free-energy landscape as a function of two parameters that are relevant to the activation process, namely the distance between the C_α_ atoms of the residues R165^3.50^ and T279^6.34^, and the root mean square deviation (RMSD) of C_α_ atoms of the NPxxYA motif (residues N332^7.49^ to A337^7.54^) to the inactive crystal structure of MOR. We also used this method to reweigh the distributions of the variables used in the n-body information theory and MaxCal analyses presented in the next sections.

### G. Information Theory Analysis

We applied the n-body information theory nBIT^13^ method to reweighted metadynamics trajectories using in-house python scripts to study the contribution of each receptor residue to the transmission of information between the ligand binding pocket and the intracellular region of the receptor. To this end, we limited our analysis to three classes of variables: (a) the first two principal components of the positions of the heavy atoms of the sidechains inside the ligand binding pocket (*PC*_1_, *PC*_2_), (b) the two CVs used to represent the activation process, namely the TM3-TM6 distance between the C_α_ atoms of residues R165^3.50^ and T279^6.34^ and the RMSD of the NPxxYA motif (NPxxYA RMSD) from the MOR inactive crystal, and (c) the Cartesian coordinates (*x, y, z*) of the C_α_ atoms of each receptor transmembrane residue.

To calculate the co-information value, we built 5-dimensional free-energy landscapes for the aforementioned parameters using the reweighting scheme^36^ described in section F. Using these free energies, we calculated the (Shannon) entropy of a set of degrees of freedom X as:

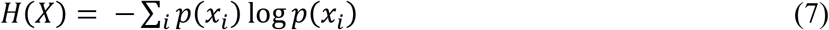

where *p*(*x*_*i*_) is the probability distribution of *x*_*i*_ ∈ *X*. The co-information can then be calculated for any set of variables as:

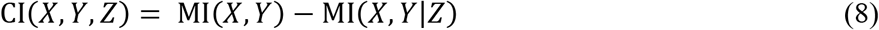

where MI is the mutual information, defined as MI(*X, Y*) = *H*(*X*) + *H*(*Y*) − *H*(*X, Y*) and the conditional mutual information and conditional entropy are defined as MI(*X*|*Z*) = *H*(*X*|*Z*) + *H*(*Y*|*Z*) − *H*(*X, Y*|*Z*) and *H*(*X*|*Z*) = *H*(*X, Z*) − *H*(*Z*), respectively. Using these definitions, it is easy to see that the 3-body co-information is fully symmetric under permutations of its three variables and that equation 8 can be recast as:

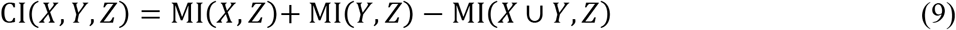

A positive value of CI(*X, Y, Z*) suggests that the sum of the information on *Z* gained from knowledge of either *X* or *Y* (i.e. MI(*X, Z*) + MI(Y, Z)) is larger than the information on *Z* gained from knowing both of *X* and *Y* (i.e. MI(*X* ∪ *Y, Z*)); hence there is redundant information pertaining to *Z* in *X* and *Y*. This can be described as a common cause structure; the correlation between *X* and *Y* is partially explained by the value of *Z*. By the same logic, a negative co-information value suggests that knowledge of both *X* and *Y* provides additional information regarding *Z*. Co-information values calculated in this way are very sensitive to the way the histograms are built and it is therefore crucial to use the same bin sizes and histograms while building the multidimensional histograms for each residue, ensuring that the ranking of the co-information values for the residues is preserved.

### H. Maximum Caliber Kinetic Model

By design, the MaxCal approach,^14^ which is the maximum entropy principle applied to dynamic quantities, allows for the construction of the most probable kinetic model that is compatible with a given free-energy landscape and with constraints that depend on the temporal evolution of the system. For a given set of stationary probabilities π_*i*_, we maximize, using in-house python scripts, the path entropy, 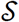, with respect to the Markov model transition probabilities *p*_*ij*_,

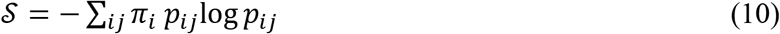

under constraints that fix the average value of dynamical quantities 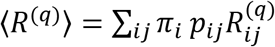 to specified values 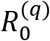. To study continuous stochastic processes, the mean jump rate 〈*N*〉 is conveniently chosen as one of these dynamical constraints. With this choice of constraints, it has been shown^14^ that the transition probabilities *p*_*ij*_ that result in maximum caliber are proportional to

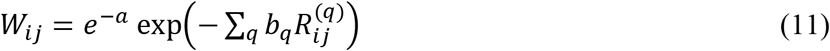

where *a* and *b*_*q*_ are the Lagrange multipliers associated with 〈*N*〉 and with all other dynamical constraints 〈*R^(*q*)^*〉, respectively. For short lag-times δ*t*, we can rewrite *W* defining a matrix Δ and making explicit the dependence on the lag-time:

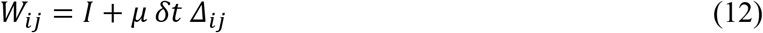

where *I* is the identity matrix, and we defined μ δ*t* = *e*^−*a*^. Further imposing the condition of detailed balance, the transition rates *κ*_ij_ can be calculated as:

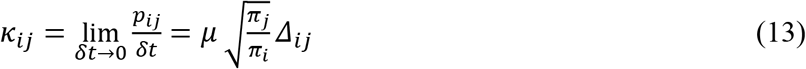

and the final transition matrix is:

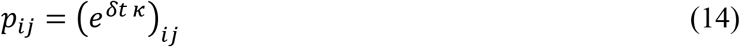

Practically, the Lagrange multipliers *a* and *b*_*q*_ are derived so that 〈*N*〉 and 〈R^(^*q*)〉 equal given values *N*_0_ and 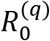 estimated from simulations or experiments.

In order to design a self-contained strategy, we introduce here a way to estimate *N*_0_ and 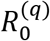 directly from the MetaD simulations. We apply concepts introduced recently for the calculation of time-lagged independent component analysis (tICA) correlation matrices^37^, and taking into consideration that the stochastic dynamics under a metadynamics bias can be rigorously described by an ordinary differential equation with established asymptotic behavior^38^. Thus, we calculate the unbiased value of the constraint averages

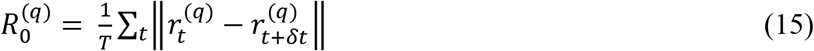

where 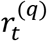 is the value of the collective variable at time *t*, and *T* is the total length of the trajectory, by reweighting the biased MetaD trajectories as^37^:

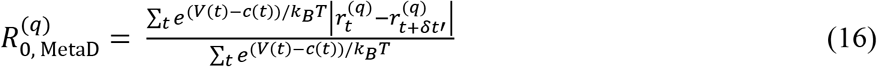

Here, the correction function *c*(*t*), which can be calculated from the bias, tends asymptotically to the irreversible work performed on the system. Since the bias accelerates the dynamics, the lag-time δ*t*′ must also be rescaled, as described in ref.^37^, so that

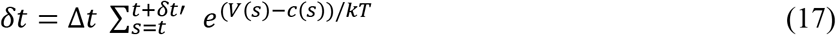

where Δ*t* is the trajectory time step. For validation purpose, the constraints calculated from the MetaD simulations were compared to those obtained from previously published HTMD adaptive sampling simulations.^11^ Specifically, using the MSMs built from the HTMD trajectories, we calculated the path ensemble averages of the distances traveled along each CV describing receptor activation (i.e., change in TM3-TM6 distance and NPxxYA RMSD from the MOR inactive crystal) per unit time, and compared these values to those obtained from MetaD by application of equation 16.

The kinetic model obtained from the transition matrix *p*_*ij*_ defined in equation 14 was then studied using standard tools for MSM analysis available in PyEMMA 2.4^39^, which yielded the mean first passage times (MFPT) reported herein.

## III. RECAPITULATING THERMODYNAMIC PROPERTIES OF GPCR ACTIVATION

As mentioned in the Introduction, our recent study of the dynamics and kinetics of the morphine-induced MOR activation process using an HTMD adaptive sampling protocol revealed two metastable regions in the free-energy landscape, referred to as intermediate I and II, in addition to metastable regions comprising crystal-like inactive and active receptor conformations. We showed in that work that these regions contain highly populated conformational states of MOR with intermediate region II exhibiting a much higher free energy compared to the other metastable regions. We proceeded to compare these results with those obtained from the MetaD-based path sampling of the morphine-induced MOR activation process (see the Computational Details section), and confirmed the finding of four main metastable regions.

The reweighted free-energy landscape obtained from the MetaD simulation of the morphine-induced activation of MOR is shown in Fig. 1(a) as a function of CVs describing major conformational changes occurring upon activation (i.e., TM3-TM6 distance and NPxxYA RMSD from the MOR inactive crystal). In this figure, overlaid gray dots correspond to the positions of the k-means centers of the free-energy landscape obtained from our previously published analysis of the HTMD adaptive sampling simulations of the morphine-induced MOR activation.^11^ As evident from this figure, the 3 μs MetaD simulations we carried out explore a free-energy landscape that is comparable to that obtained by ∼240 μs HTMD adaptive sampling simulations, with the only exception of regions with very high free energies (i.e., intermediate region II). We further calculated the populations of each metastable region, i.e., the crystal-like inactive and active, intermediate I, and intermediate II regions of the free-energy landscape of the morphine-induced MOR activation, and found good agreement between the two approaches as shown in Fig. 1(b). Taken together, these results suggest that well-tempered MetaD using path CVs is capable of recapitulating the thermodynamic properties of complex processes, such as GPCR activation, as seen in more computationally intensive methods such as the HTMD adaptive sampling protocol.

**Figure 1.**
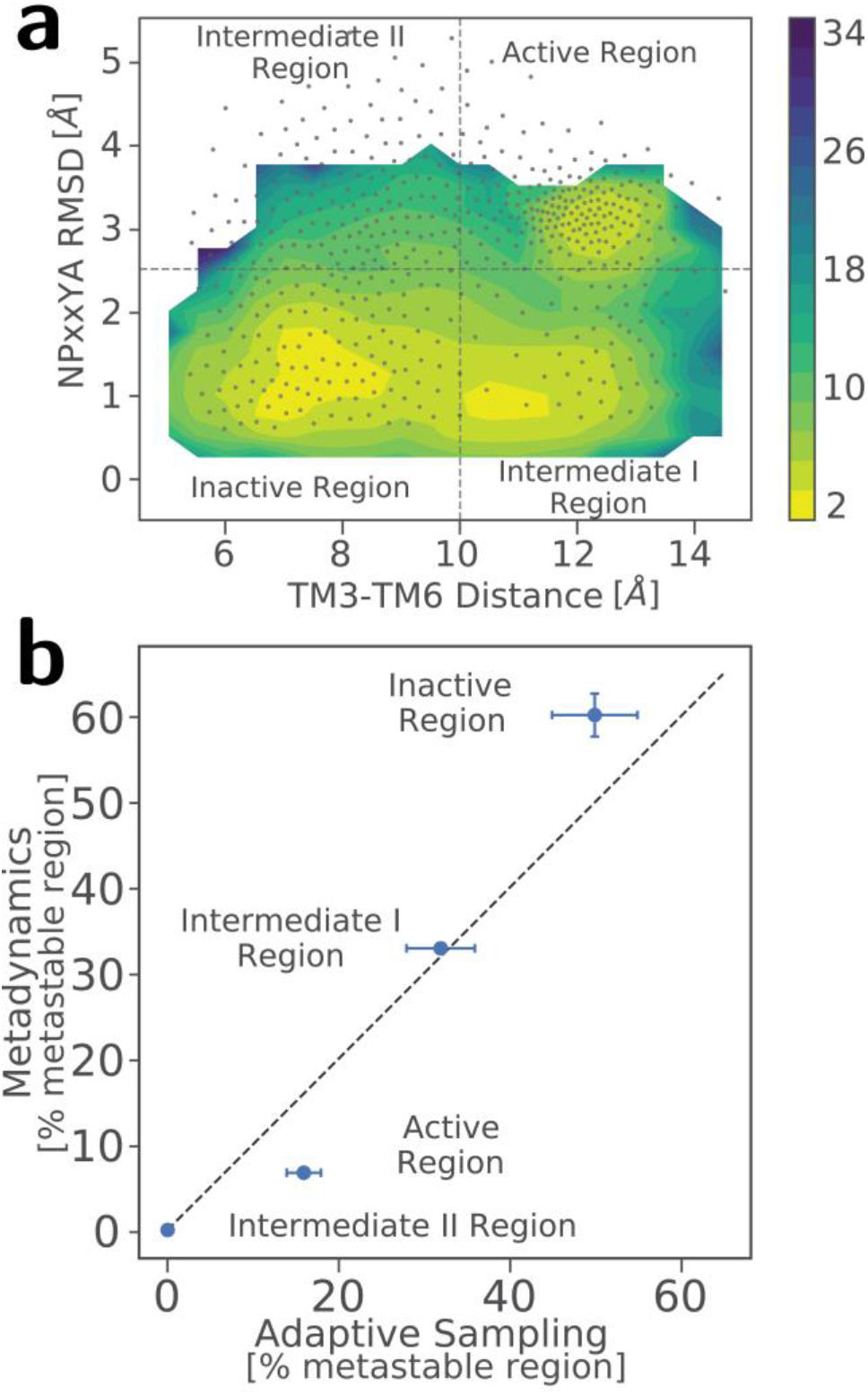
Comparison between thermodynamic properties from MetaD and the HTMD adaptive sampling protocol. (a) Free-energy landscape of the MetaD simulations of the morphine-bound MOR as a function of CVs describing major conformational changes upon receptor activation (i.e., TM3-TM6 distance and NPxxYA RMSD from the MOR inactive crystal). Overlaid gray dots refer to the positions of the k-means centers of the free-energy landscape obtained from our previously published HTMD simulations of the morphine-induced MOR activation. (b) Correlation between the populations of each identified metastable region, i.e., the crystal-like active and inactive, intermediate I, and intermediate II regions, of the free-energy landscape of the morphine-induced MOR activation derived from MetaD or HTMD simulations.

## IV. REPLICATING INFORMATION TRANSFER ACROSS THE RECEPTOR

In addition to replicating free-energy landscapes, it is also important to verify that molecular dynamic details obtained from the MetaD-based strategy proposed herein can be used to capture how information is transferred from the ligand-binding pocket to the intracellular G protein-binding region of the receptor. To this end, we calculated (see the Computational Details section) the contribution of each receptor residue to the mutual information between the ligand binding pocket and the intracellular region of the receptor upon activation. This contribution is quantified by a three-body co-information value calculated using (a) the first two principal components of the positions of the heavy atoms of the sidechains inside the ligand binding pocket, (b) the two CVs used to represent the activation process, namely the TM3-TM6 distance and the NPxxYA RMSD from the MOR inactive crystal, and (c) the Cartesian coordinates of the C_α_ atoms of each receptor transmembrane residue. Co-information values larger than 1 (in absolute value) are listed in Supplementary Table I.

The calculated co-information values from the MetaD and HTMD simulation trajectories are compared in Fig. 2(a). As mentioned in the Computational Details section, the more negative a co-information value is for a given residue, the more significant is that residue’s contribution to the information transfer between the extra- and intra-cellular regions of the receptor. A visualization of the location of the most contributing residues to the information transfer in MOR as calculated from MetaD or HTMD simulations is provided in Fig. 2(b) and 2(c), respectively. As can be seen in these figures, the two simulation protocols yield similar co-information patterns, with the majority of highly contributing residues to the information transfer (Supplementary Table I and red color in Fig. 2(b) and 2(c)) either located in the lower halves of TM5, 6, and 7 and in H8, or in the extracellular regions of TM1, 6, and 7. These results suggest that the well-tempered MetaD strategy reported herein replicates the dynamical information obtained by HTMD simulations, and can therefore be employed as an effective tool for studying allostery in GPCRs.

**Figure 2.**
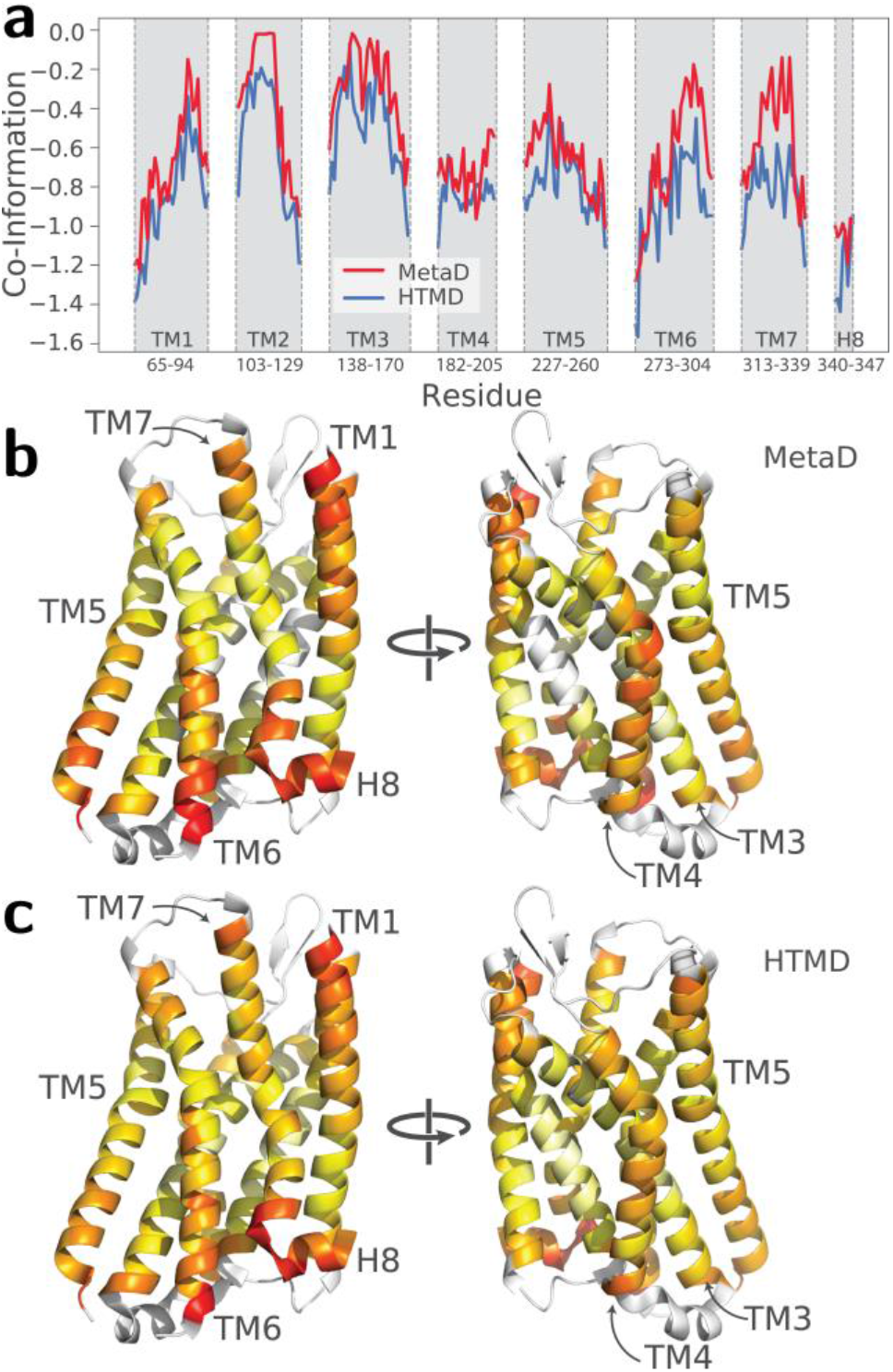
Comparison between co-information values derived from MetaD and HTMD simulations. (a) Normalized co-information values from MetaD and HTMD simulations plotted for each MOR transmembrane residue. (b, c) Normalized co-information values depicted on the inactive crystal structure of MOR using a color scheme ranging from white to yellow to red, where red represents the highest contributing residues to the information transfer between the ligand binding pocket and the intracellular region of MOR, as derived from MetaD and HTMD simulations, respectively. Co-information values for the loop regions were not calculated.

## V. DERIVING KINETIC PROPERTIES OF GPCR ACTIVATION FROM METAD USING MAXCAL PRINCIPLE

Deriving kinetic rates directly from standard MetaD simulations is not possible because transitions are accelerated by a bias potential in these simulations. The recently proposed infrequent metadynamics^40, 41^ strategy based on transition state theory^40^ allows to extract unbiased kinetic information from biased trajectories, but it requires a reduced bias deposition rate, as well as multiple validation simulations. Another recently proposed strategy based on Girsanov reweighting of the transition matrix^42^ is promising, but has yet to be validated for large biological systems. Here, we test the efficiency and accuracy of the MaxCal principle in extracting kinetic rates from MetaD. Specifically, the MaxCal principle, which is the maximum entropy principle applied to dynamic quantities, allows one to construct the most probable, minimally biased kinetic model that is compatible with a given stationary distribution of the system and with a number of constraints imposed on the system’s kinetics that are derived here from MetaD.^14, 16, 43^

Although this approach has previously been tested on various small peptides,^14, 43^ GPCRs are far more complex systems and an assessment that MaxCal could accurately capture the kinetics of GPCR activation was necessary prior to its application to MetaD-derived free energies. Thus, we first applied MaxCal to the free-energy landscape of morphine-induced MOR activation derived from the more computationally expensive HTMD adaptive sampling protocol^11^. To capture the system’s dynamics as a function of two commonly used variables that describe the activation process, we constrained the path ensemble averages of (a) the difference in TM3-TM6 distance between pairs of microstates, and (b) the difference in NPxxYA RMSD from the MOR inactive crystal structure between pairs of microstates. To compare the kinetic model derived from MaxCal to that obtained from the direct MSM analysis of the HTMD trajectories, we calculated the path ensemble averages to be used as constraints from the MSM built from these trajectories (see Supplementary Table II), and then applied them during path entropy maximization. Despite using only two constraints, the MFPTs between the four metastable regions of the free-energy landscape of morphine-induced MOR activation derived from the HTMD- and the MaxCal-derived MSMs (Supplementary Fig. 2(a) and 2(b), respectively) are in excellent agreement as shown by their correlation in Supplementary Fig. 2(c). This observation suggests that MaxCal is capable of replicating the kinetics of GPCR activation estimated from MSM analysis of the HTMD simulations.

To assess the effectiveness of MaxCal in obtaining from MetaD MOR activation kinetics comparable to that obtained from the HTMD adaptive sampling protocol,^11^ we recalculated the values of the aforementioned path ensemble averages from the MetaD simulations. These values are in good agreement with those obtained from the HTMD MSMs (see Table II in the SI). These MetaD-derived averages were then used, together with the MetaD-derived free energies, to constrain the path entropy during its maximization. The MFPTs between the most populated metastable states obtained from the HTMD-derived MSMs and those obtained by MaxCal from the MetaD runs, are shown in Fig. 3(a) and 3(b), respectively. As shown in this figure, the MFPTs from MetaD simulations are, on average, within a factor of 2 of the corresponding HTMD adaptive sampling results. In contrast, the MaxCal approach failed to replicate the kinetic behavior of the system’s intermediate region II, which contains very high free-energy areas at NPxxYA RMSDs larger than 4 Å.

**Figure 3.**
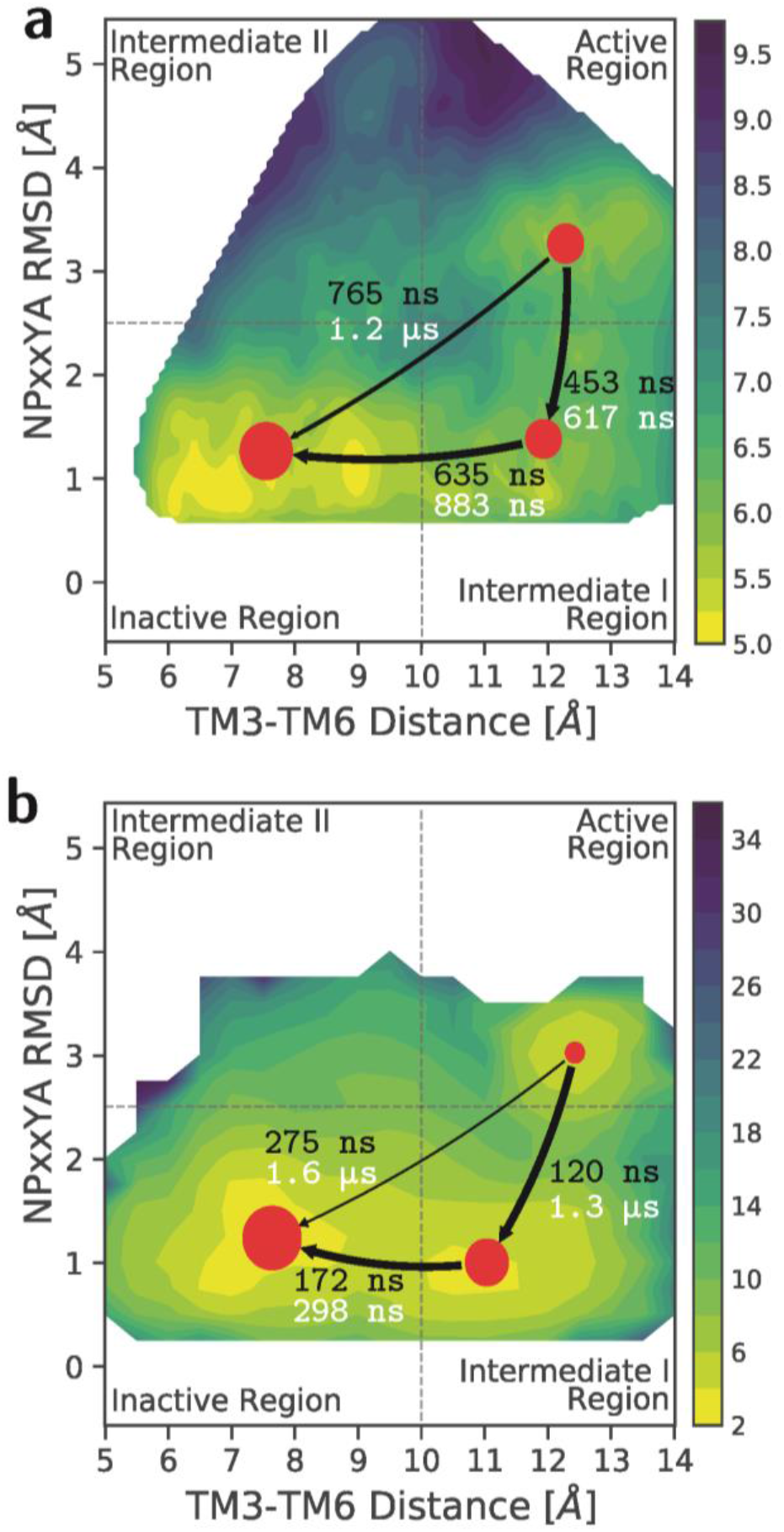
Comparison between kinetic properties of MOR activation derived from MetaD and HTMD simulations. The Mean first passage times (MFPTs) obtained for the most populated regions of the free-energy landscapes of morphine-induced MOR activation from (a) the MSMs built using the HTMD adaptive sampling runs and (b) the MaxCal MSMs built using the MetaD runs.13

Given the difficulty associated with extracting kinetic rates from standard MetaD simulations, it is gratifying that the MaxCal approach provides a means for successfully estimating kinetic rates for metastable regions that are most relevant to GPCR activation. We propose that additional constraints derived experimentally may further increase the accuracy of kinetic properties obtained by this integrated MetaD-MaxCal strategy.

## VI. SUMMARY AND CONCLUSIONS

Our results confirm that well-tempered MetaD with path collective variables is a fast and reliable method for studying the thermodynamic properties of GPCR activation. Furthermore, the propagation of information across the receptor calculated from MetaD simulations is comparable to that derived from adaptive sampling protocols, suggesting that this simulation method also offers a more efficient strategy for studying allostery in GPCRs. Finally, we show that kinetic properties of complex systems can be derived from MetaD simulations using the MaxCal principle, and these results are comparable to those obtained from more computationally expensive adaptive sampling protocols.

In conclusion, MetaD can be employed as an efficient method for studying ligand-induced activation mechanisms in GPCRs by reducing the necessary computation time by ∼2 orders of magnitude, and its combination with MaxCal allows to derive kinetic properties that are in agreement with those obtained by computationally more expensive methods.

## ACKNOWLEDGMENTS

This work was supported by National Institutes of Health grants DA026434 and DA034049. Computations were run on resources available through the Scientific Computing Facility at Mount Sinai and the Extreme Science and Engineering Discovery Environment under MCB080077, which is supported by National Science Foundation grant number ACI-1053575.

